# Bacteriophage adaptation to a mammalian mucosa reveals a trans-domain evolutionary axis

**DOI:** 10.1101/2021.05.11.443681

**Authors:** Wai Hoe Chin, Ciaren Kett, Oren Cooper, Deike Müseler, Yaqi Zhang, Rebecca Bamert, Ruzeen Patwa, Laura C. Woods, Citsabehsan Devendran, Joe Tiralongo, Trevor Lithgow, Mike J. McDonald, Adrian Neild, Jeremy J. Barr

## Abstract

The majority of viruses within the human gut are obligate bacterial viruses known as bacteriophages (phages)^1^. Their bacteriotropism underscores the study of phage ecology in the gut, where they sustain top-down control^2–4^ and co-evolve^5^ with gut bacterial communities. Traditionally, these were investigated empirically via *in vitro* experimental evolution^6–8^ and more recently, *in vivo* models were adopted to account for gut niche effects^4,9^. Here, we probed beyond conventional phage-bacteria co-evolution to investigate the potential evolutionary interactions between phages and the mammalian “host”. To capture the role of the mammalian host, we recapitulated a life-like mammalian gut mucosa using *in vitro* lab-on-a-chip devices (to wit, the gut-on-a-chip) and showed that the mucosal environment supports stable phage-bacteria co-existence. Next, we experimentally evolved phage populations within the gut-on-a-chip devices and discovered that phages adapt by *de novo* mutations and genetic recombination. We found that a single mutation in the phage capsid protein Hoc – known to facilitate phage adherence to mucus^10^ – caused altered phage binding to fucosylated mucin glycans. We demonstrated that the altered glycan-binding phenotype provided the evolved mutant phage a competitive fitness advantage over their ancestral wildtype phage in the gut-on-a-chip mucosal environment. Collectively, our findings revealed that phages – in addition to their evolutionary relationship with bacteria – are also able to engage in evolution with the mammalian host.

## Introduction

Bacteriophages (phages) are viruses that predate bacteria to replicate. Their bacteriotropism is reflected by the manifold of studies on phage-bacteria antagonistic co-evolution^6–8,11^. In the mammalian gut, this antagonistic co-evolutionary dynamic is key in maintaining long-term phage-microbiome homeostasis and diversity^9,12^. While extremely insightful, this phage-to-bacteria focus has overlooked another potential trans-domain evolution: the phage-mammalian axis. Phages have been demonstrated to adhere directly to mammalian mucin^10^, and when applied to mucosal layers, phages can exhibit enhanced virulence towards bacterial hosts^13–16^. At an ecological level, the gut mucosa segregates phage and bacterial populations, establishing spatial refuges, which can promote phage-bacteria co-existence^17^. We reasoned that the direct interaction between phages and the gut mucosa could have far-reaching implications for phage persistence, ecology, and evolution. We hypothesised that phages not only engage in antagonistic co-evolution with their bacterial hosts^6^, but also evolve in response to their mammalian host or mammalian-derived factors. Using an *in vitro* labon-chip device to simulate a life-like mammalian mucosal layer^18^, we tested if phage evolution would favour phenotypes that persist within the mammalian mucosal environment.

### The gut-on-a-chip supports phage-bacteria co-existence within a mucosal layer

To investigate the capacity of phages to adapt to the mammalian mucosal environment, we fabricated a simple gut-on-a-chip microfluidic device that recapitulates key features of the mammalian gut mucosa (Fig.1A). These devices are experimentally amenable, provide an accessible platform for biological replication, and recapitulate essential organ-level functions of the gut^18,19^ (Fig.1B). Our gut-on-a-chip consisted of a single channel containing a confluent colonic cell layer, capable of mucus secretion and exhibiting mucus turnover (Fig.1C, Supplementary Fig.1A). The gut-on-a-chip was able to support stable phage-bacteria co-existence for up to 24 hours. In each device, *Escherichia coli* bacteria and T4 phages were infused and the co-culture was maintained for 24 hours under constant perfusion with sterile media. To assess microbial population dynamics, an automated sample collection system was developed where egressing samples from the device were heat-inactivated, followed by collection at 30-minute intervals, with phages and bacteria subsequently quantified via quantitative PCR (qPCR) (Fig.1E, Supplementary Fig.1B). Here, phage numbers increased rapidly within two hours post-inoculation with a concomitant crash in the bacterial population. Following this crash, the phage population stabilised between 10^6^ – 10^7^ phage/ml whilst maintaining suppression of the bacterial population, particularly over the first 10 hours. Subsequently, bacterial levels rose above qPCR detection thresholds, exhibiting classical prey-predator dynamics, characterised by cyclic changes in phage-bacterial numbers^20^. There was considerable variation between replicate devices in both phage-bacteria population abundances and dynamics, suggesting that each device was delineated by inherent fluctuations and ecological stochasticity. This was exemplified in one replicate (Fig. 1E, replicate 1) where the bacterial population remained suppressed below qPCR detection threshold while the remaining replicates (Fig.1E, replicates 2 and 3) exhibited detectable but disparate phage-bacterial population dynamics overtime. Overall, the gut-on-a-chip provided a tripartite model system that supported mammalian, bacterial and phage co-culture.

**Fig.1.**
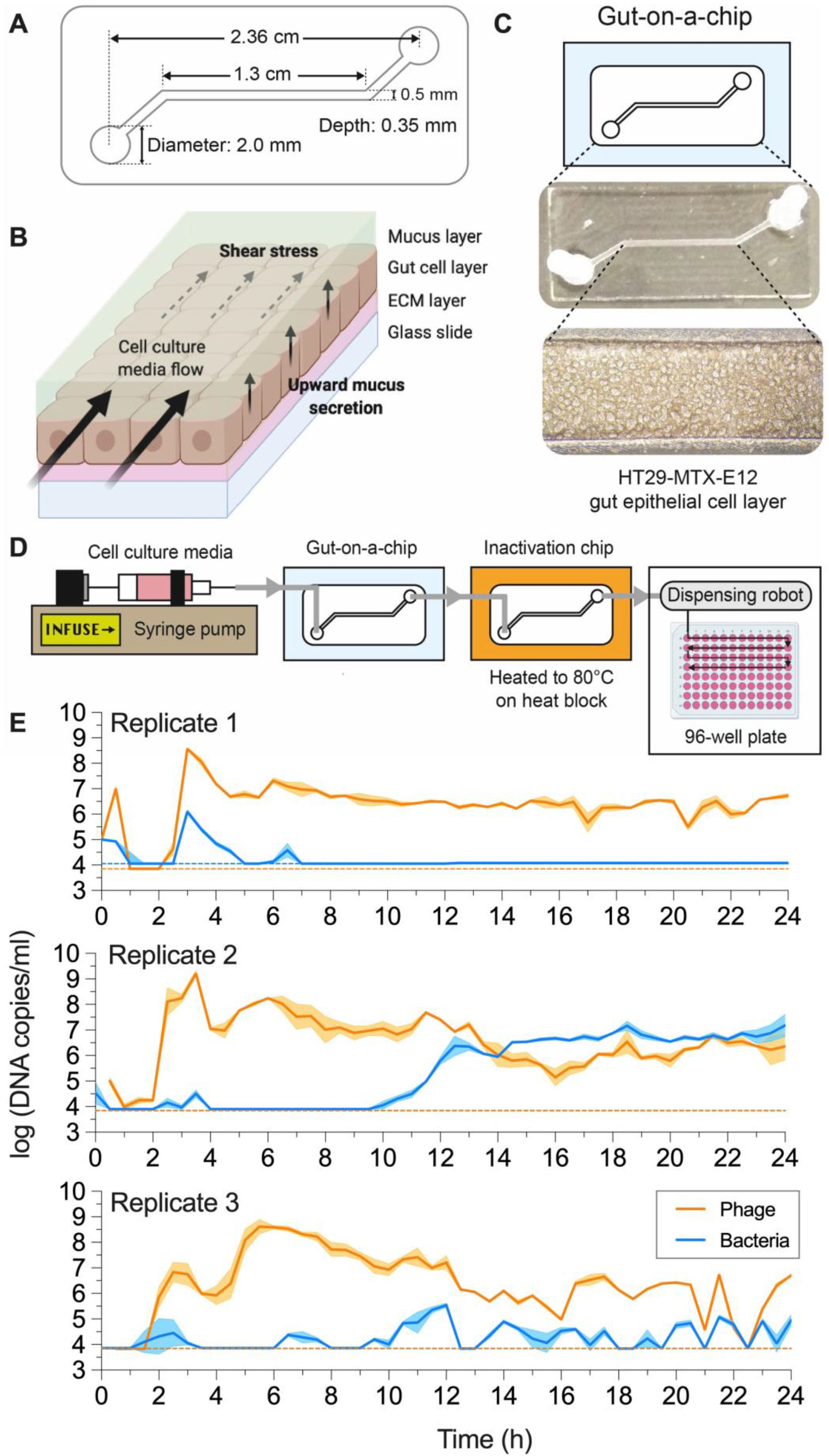
The gut-on-a-chip mucus environment supports phage-bacteria co-existence. A) Schematic and channel dimension of the gut-on-a-chip. B) Mucus turnover dynamics from the device is driven by shear stress from fluid flow and upward mucus secretion from the epithelial layer. C) HT29-MTX-E12 cell line grows and differentiates within the device channel environment to produce a mucus layer at ~72 hours post-seeding. D) Schematic for overall gut-on-a-chip set-up for chip perfusion, continuous sample inactivation via heat and automated sample collection for qPCR quantification. E) qPCR-quantified phage-bacteria population in three separate devices at 30-minute intervals over 24 hours. Plotted line and shaded region represents mean ± SEM of three qPCR technical replicates (n = 3) per experimental replicate (N = 3). Orange and blue dotted lines represent the qPCR limit of detection threshold for phage and bacteria respectively, per biological replicate.

### The mammalian mucus layer influences phage evolution

We performed experimental evolution of phage populations within the mucosal environment of the gut-on-a-chip (Fig.2A). Gut-on-a-chip devices were inoculated with populations of T4 phages and *E. coli* bacteria (the founding phage population herein referred to as the “ancestral” phage), which were maintained for 24 hours. We then conducted successive transfers of the evolved phage populations into fresh gut-on-a-chip devices grown from naïve gut cells and seeded with naïve bacterial populations (“naïve” referring to entities that had no prior exposure to phages). By limiting the transfers to the phage population only, we directed phage adaptation towards the mucosal environment, while limiting phage-bacterial co-evolution. In total, we performed five successive transfers of evolved phages across three biological replicates in gut-on-a-chip devices and in test-tubes; the latter as an experimental control lacking a mammalian mucosal environment. Phages and bacteria were consistently recovered from the gut-on-a-chip mucosal environments, demonstrating that the mucus layer supported phage propagation while maintaining a stable bacterial population over the course of the experimental evolution (Fig.2B, gut-on-a-chip). This contrasted with controls populations in test-tubes where bacteria were frequently extinct (Fig.2B, test-tube).

**Fig.2.**
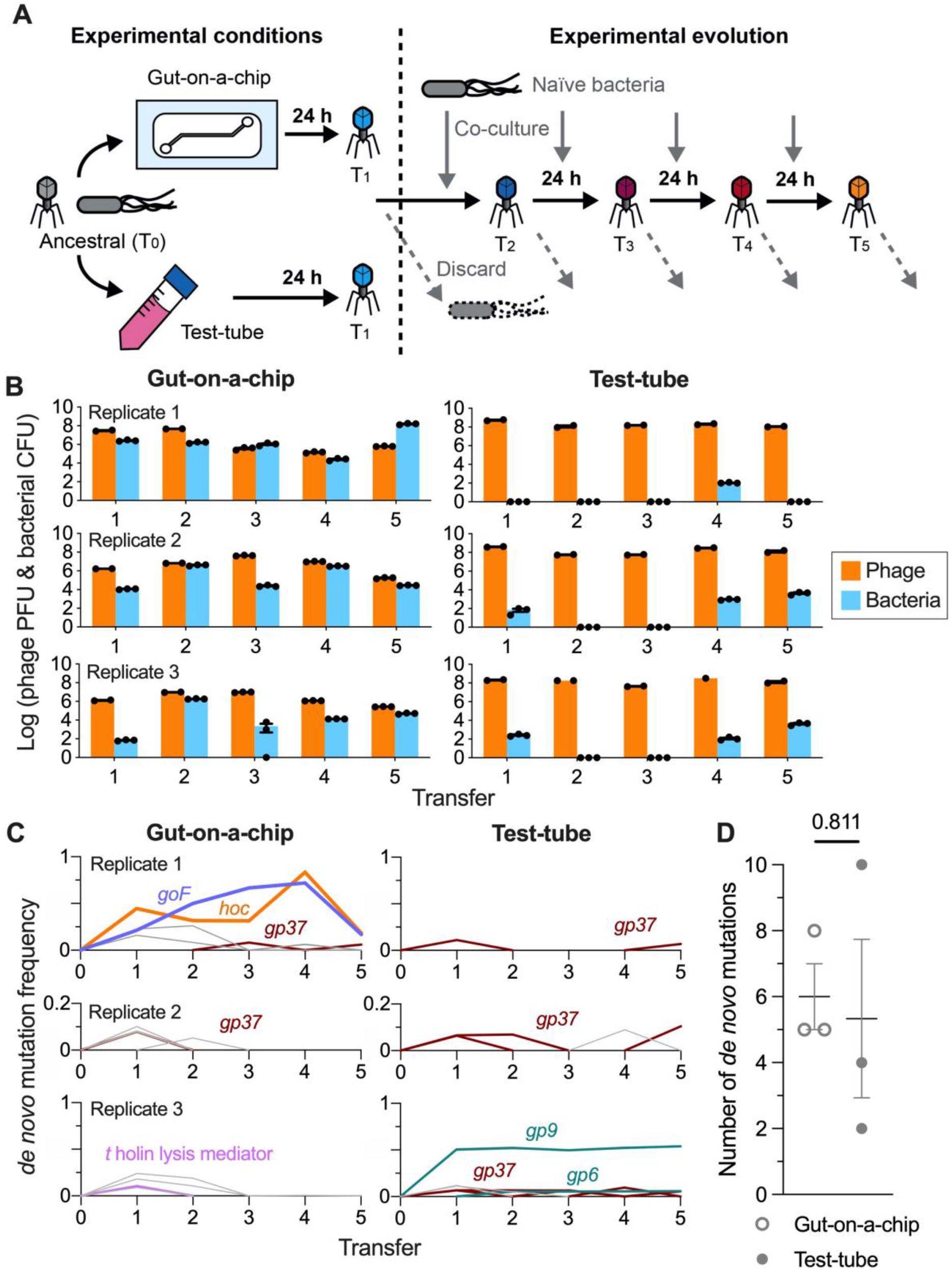
Phages evolve in response to the life-like mammalian mucus layer in the gut-on-a-chip. A) Ancestral (zeroth transfer; T_0_) T4 phage and *E. coli* bacterial hosts were inoculated into three gut-on-a-chip and test-tube set-ups, respectively. The co-cultures were incubated for 24 hours with phages subsequently harvested for the first transfer (T_1_). Phages from T_1_ were transferred onto fresh chips and test-tubes seeded with naïve *E. coli* B hosts and the process was repeated till the fifth transfer (T_5_). B) Population of phages and bacteria from the mucus sample at the end of each 24-hour passage in chip and test-tube replicates. C) Frequency of *de novo* mutations emerging from the phage population over five transfers from the gut-on-a-chip and test-tube set-ups. Coloured line represents the mutations: D246N *hoc* mutation in orange, Δ21bp *goF* mutation in purple, *gp37* (distal subunit phage long tail fibre) in brown, *t* holin lysis mediator in pink, and *gp6* and *gp9* (phage baseplate subunits) in teal. Grey lines represent other transient and low-frequency *de novo* mutations (see Supplementary table 1B). D) Average number of *de novo* mutations from phage populations evolved in gut-on-a-chip and test-tube conditions. Data points in panels B were technical replicates for phage-bacteria quantification from transfers, while datapoints in D were independent experimental replicates with values plotted as mean ± SEM across the experimental replicates (N = 3). P-value in panel D was derived from a two-tailed unpaired t-test.

Next, we sought to determine the evolutionary changes that occurred in the phage populations between gut-on-a-chip and test-tubes using whole-genome sequencing, followed by read alignment and mutational calling. We discovered background mutations comprising of single nucleotide polymorphisms (SNPs) and singl-enucleotide insertions in our ancestral phage population, reflective of their long-term laboratory storage and genetic drift^21^ (Supplementary table 1A). We subtracted these background SNPs and insertions from our mutational readouts in order to highlight *de novo* mutations in our gut-on-a-chip and test-tube phage populations.

In the case of our gut-on-a-chip populations, *de novo* mutations were found in genes encoding nucleotide binding and metabolism proteins, structural proteins and hypothetical proteins, most of which were transient and low abundance (Supplementary Table 1B). While we did not observe parallel evolution across our chip-evolved populations, two mutations attained high-abundance within the first replicate population. The first was a non-synonymous SNP within the *hoc* (highly immunogenic outer capsid) gene, which encodes for an accessory outer capsid protein that has been demonstrated to facilitate phage adherence to mucus^10^. This SNP resulted in an amino acid change at position 246 from aspartic acid to asparagine (henceforth referred to as D246N Hoc). The second was an in-frame 21bp-deletion (Δ21bp) of the *goF* gene which encodes for a transcription antitermination factor that antagonises the bacterial ρ (Rho) termination factor from prematurely degrading phage mRNA transcripts (Supplementary Fig.2A & 2B)^22^. At their peak frequencies in the fourth transfer, both Δ21bp *goF* and D246N Hoc mutations achieved 72% and 83.3% of the population respectively, indicating a strong selective advantage for these mutations within the gut-on-a-chip (Fig.2C, gut-on-a-chip replicate 1).

By contrast, phages evolving in test-tubes exhibited arms-race-like dynamics with their bacterial hosts; meaning that mutations found were largely directed towards adaptation for bacterial infection^6,11^. The phages evolved in test-tubes exhibited *de novo* mutations in *gp37*, which encodes the distal subunit of the phage long tail fibre responsible for phage adsorption onto its bacterial hosts. These mutations were observed transiently and at low-frequency across all test-tube replicate populations (Fig.2C, test-tube replicates; Supplementary Table 1A). We also observed other mutations affecting phage baseplate-associated genes (*gp6* and *gp9*), whose gene products facilitate genome injection into the bacterial host during infection; although these mutations were only present in one replicate (Fig.2C, test-tube replicate 3). Despite the disparate *de novo* mutation profiles between gut-on-a-chip and test-tube populations, we did not observe significant differences in total number of *de novo* mutations across the five transfers between the gut-on-a-chip (6.3 ± 0.9 mutations) and test-tube phage populations (7.0 ± 2.1 mutations) (Fig.2D).

### High multiplicity-of-infection is a driver for phage recombination

In asexual populations such as with phages, genetic recombination is key to enhancing fitness by alleviating clonal interference and genetic hitch-hiking^23,24^. For lytic phages, such as T4, recombination occurs when multiple phage genotypes co-infect the same bacterial host, allowing allelic exchange between the phage genomes^25^. Since co-infections drive recombination, higher multiplicity-of-infections (MOIs) typically render higher recombination rates^25^. Crucially, high MOIs were sustained in our chip-evolved phage populations, where elevated phage-to-bacteria ratios were observed (Fig.1E & 2B). We noted that the D246N Hoc and Δ21bp *goF* mutations follow similar frequency trajectories with their increase and decline between the fourth and fifth transfers (Fig.2C, test-tube replicate 1). Their intertwined trajectories surpassing 50% frequencies suggest that the mutations had recombined onto a shared genetic background to overcome clonal interference. We sought to verify if the high phage-to-bacteria ratios – and thus, high MOI – were drivers for recombination in lytic phage populations. We initiated one-step phage growth experiments at high and low MOIs (i.e. 10 and 0.1 respectively) with a 1:1 mix of two phage mutants: i) experimentally-derived Δ21bp *goF* mutant (gene position: 5842 – 6267) and ii) lab-stock *hoc* deletion mutant (Δ*hoc*; gene position: 110187 – 111317) (Fig.3A). By limiting the phages to a single growth step, we limit phage recombination within a single replicative cycle. Following PCR screening of individual plaques, we found that 44% of phage progeny were recombinants at high MOI conditions, with a bias towards wildtype recombinants (43/98 phages screened were recombinants; 31/43 of wildtype recombinants; Fig.3B, Supplementary Fig.3). Meanwhile, only 1 wildtype recombinant phage was detected from 98 isolates screened from low MOI conditions, i.e. ~1% recombinant frequency (Fig.3B, Supplementary Fig.3). The rapid emergence of recombinants within a single phage replication cycle suggests that recombination is a key driving force for phage evolution. Collectively, this suggests that a high and sustained phage-to-bacteria ratio facilitates genetic recombination in phages, which in-turn promote selection of high-frequency beneficial mutations and alleviation of clonal interference.

**Fig.3.**
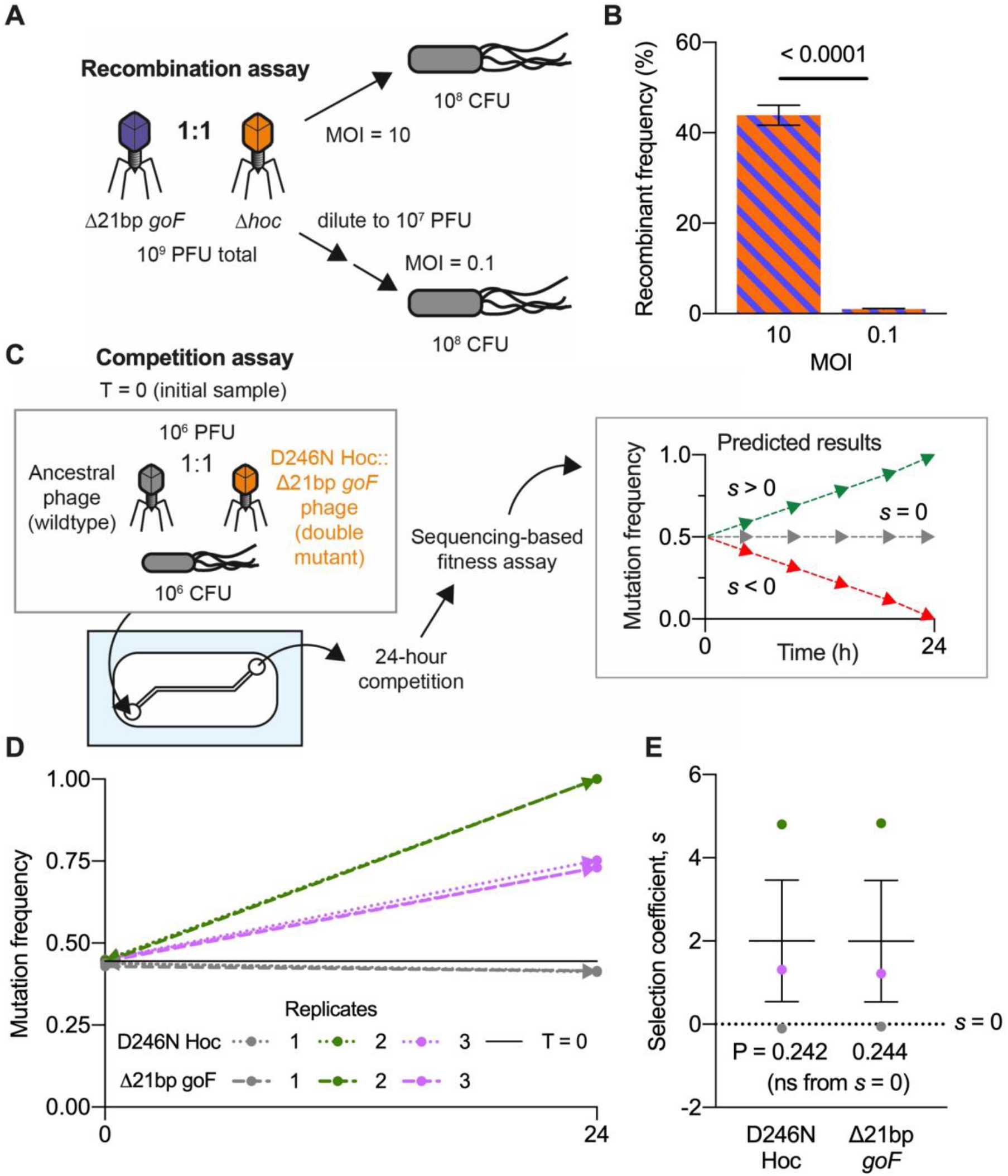
Mucus layer supports phage recombination and selection of beneficial mutants. A) Gut-on-a-chip-evolved Δ21bp *goF* phage mutant was mixed at a 1:1 ratio with lab-derived *Δhoc* mutant. Naïve *E. coli* were infected with the 1:1 phage mixture at MOIs 10 and 0.1 following a one-step growth protocol to ensure that only a single round of viral replication could occur. B) Percentage frequency of phage recombinants from PCR screening for Δ21bp *goF-Δhoc* or wildtype-reconstituted recombinants. 49 phage isolates (n = 49) were screened per experimental replicate (N = 2) leading to a total of 98 isolates screened per experimental condition (MOI 10 or 0.1) (Supplementary Fig.3). C) Competition experiment between D246N Hoc::Δ21bp-*goF* mutant phage with ancestral phage T4 in the gut-on-a-chip. Gut-on-a-chip seeded with naïve *E. coli* was inoculated with equal proportions of the respective phage genotypes. Chip effluents collected at timepoints T = 0 and 24 hours were subjected to whole-genome sequencing to track D246N Hoc and Δ21bp-*goF* mutations after 24 hours of competition. Estimated selection coefficients could be positive (*s* > 0), neutral (*s* = 0) or negative (*s* < 0). D) D246N Hoc and Δ21bp *goF* mutational frequencies measured from three independent gut-on-a-chip replicates (N = 3) between T = 0 and 24 hours. E) Plot of estimated mean selection coefficient for D246N Hoc and Δ21bp *goF* mutation in each experimental replicate (N = 3). Black solid line in panel D represents the initial (T = 0) average frequency of D246N Hoc mutation at 44.5% and Δ21bp *goF* mutation at 43.9%, across three replicates (N = 3). Error bars in panel B and line with error bars in panel E represent mean ± SEM across experimental replicates. P-values in panel B were derived from unpaired t-test between treatment conditions (MOI 10 and 0.1) and; in panel E, between coefficients D246N Hoc and Δ21bp *goF* mutations against *s* = 0 (no selection).

### Phage mutant outcompetes ancestor phage in mucus

To assess the fitness of the evolved phage possessing the D246N Hoc and Δ21bp *goF* mutations, we competed the evolved double mutant phage against its ancestral counterpart in the gut-on-a-chip mucus environment. The double mutant phage was isolated and genotypically verified through Sanger sequencing. Competition between the double mutant and ancestral phage was initiated by inoculating both phages at a 1:1 ratio into a gut-on-a-chip, seeded with naïve bacterial host. The device effluent was sampled at 0 and 24 hours, with samples subsequently whole-genome sequenced to track the D246N Hoc::Δ21bp-*goF* double mutant frequency over 24 hours of competition (Fig.3C). We verified that our devices were accurately seeded with roughly equal proportions of mutant and wildtype phages as reflected by ~44% frequency of both the D246N Hoc and Δ21bp *goF* mutations at the initial experimental timepoint (t = 0) (Fig.3D). We observed the double mutant out-competed the wildtype phage in two of three replicate devices, eventually fixing in one of the replicate populations, while the remaining replicate showed no change from initial frequency (Fig.3D). To ascertain the strength of selection, we quantified the selection coefficients (*s*) across the replicate populations with coefficients being either positive (*s* > 0), neutral (*s* = 0) or negative (*s* < 0) (Fig.3C, Supplementary table 2). Overall, we found positive selection with *s* = 2 on average, for both D246N Hoc and Δ21bp *goF* genotypes within the mucus environment, although significance from null selection i.e. *s* = 0, was not attained due to significant variability between replicate measurements (Fig.3E, Supplementary table 2; coefficients reported as mean ± SEM with P-values derived from unpaired t-test).

### Hoc mutation alters phage mucus-adherence phenotype

Phage adherence to mucus has been described as a mechanism that facilitates phage enrichment and persistence within the mammalian mucosal layers^10^. For T4 phage, this adherence phenotype is facilitated by the outer capsid protein Hoc, which has three externally-displayed immunoglobulin (Ig)-like domains and a highly-conserved fourth C-terminal capsid-binding domain^26,27^. The D246N Hoc mutation removes an acidic residue (aspartic acid) and replaces it with a neutral residue (asparagine). This mutation is located within the third Ig-like domain, potentially altering Hoc binding affinity to mucin glycans (Fig.4A). To test for altered glycan adherence, we fluorescently labelled whole phage particles of wildtype Hoc, D246N Hoc, and *Δhoc* genotypes, and assayed for glycan binding on a microarray printed with 153 unique glycan structures (Supplementary Table 3). Binding was measured as fold-changes relative to the array background signal and verified for P-value significance. Overall, we were able to observe binding of whole phages across seven glycan families. D246N Hoc phages generally exhibited altered glycan-binding compared to wildtype phage, while Δ*hoc* phages had lower overall fold-change intensities relative to wildtype and D246N Hoc phages (Fig.4B, whole phages). To further investigate the specificity of Hoc-glycan interactions, we recombinantly-expressed wildtype and D246N Hoc proteins (Supplementary Fig.4) and tested the proteins on the glycan array. We showed that the Hoc protein-glycan binding largely matched whole phage binding results (Fig.4B, recombinant Hoc protein). Next, surface plasmon resonance (SPR) was adopted to quantify the binding strength between glycans and surface-immobilised Hoc protein. We focused on a subset of 26 glycans that were amenable for SPR measurements taken in solution under flow (see Supplementary Table 3 for full glycan array analysis). The SPR data demonstrated that both wildtype and D246N Hoc-glycan binding was specific for interactions with the same subset of fucosylated glycans (Fig.4B, Hoc-glycan SPR). Furthermore, the D246N Hoc protein had higher dissociation values (*K_D_*), indicating weaker binding to the subset of fucosylated glycans than the wildtype Hoc (Fig.4B, Hoc-glycan SPR; Supplementary Fig.5). Fucosylated mucin glycans are ubiquitous along the human gastrointestinal tract in individuals possessing a functional copy of the α-1,2-fucosyltransferase (*FUT2*) gene (known as “secretors”)^28^. Our gut-on-a-chip HT29-MTX-E12 cell line possesses *FUT2* and is capable of producing a fucosylated mucus layer in-line with the “secretor” phenotype^29^. Collectively, our results revealing Hoc-specific binding to fucosylated glycans, coupled with changes in glycan-binding affinity, indicate a direct adaptation of T4 phage to the gut-on-a-chip mucus layer.

**Fig.4:**
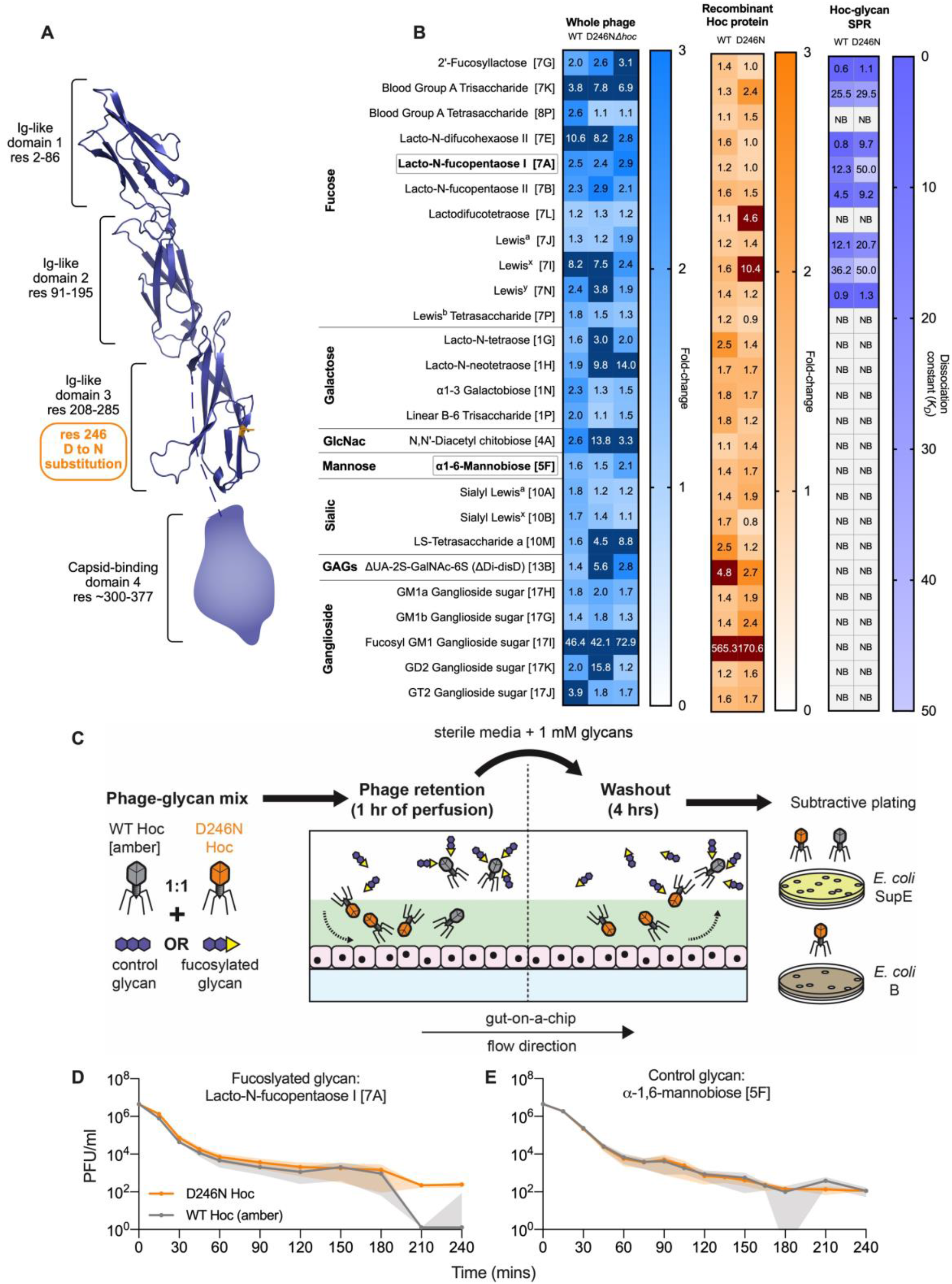
Phage evolved in mammalian mucus layer exhibit altered mucus adherence phenotype. A) T4 Hoc protein structure model demonstrating the position of D246N mutation within the third Ig-like domain, highlighted in orange. The capsid-binding fourth domain was not modelled due to the lack of structural homologues in Protein Data Bank (PDB). B) Normalised fold-change fluorescence intensities of 26 top glycan array hits (glycan ID corresponding to Supplementary Table 3 as indicated in square brackets) of labelled, ultrapurified whole phages: wildtype [WT], D246N and Δ*hoc* – blue heatmap; and recombinantly expressed Hoc proteins: WT and D246N – orange heatmap; followed by SPR assessing glycan-to-Hoc protein binding strength – purple heatmap. Numerical values in glycan array heatmaps represent fold-change magnitude normalised against background fluorescence where dark-colour panels indicate high fold-change values that were out-of-bounds from heatmap gradient. Numerical values in SPR heatmap represent dissociation constant (*K_D_*) values where higher *K_D_* values indicate lower binding affinity. “NB” in SPR heatmap indicates no binding event. Encircled and bolded glycans 7A (Lacto-N-fucopentaose I) and 5F (α-1,6-mannobiose) represent the glycans selected for phage retention and washout experiments in panels D and E. C) Experimental set-up for phage retention and washout from the gut-on-a-chip, where equal proportions of WT Hoc (with *am43^-^/44^-^* mutation) and D246N Hoc phages in 1 mM glycan solutions were perfused in the gut-on-a-chip for an hour during the retention phase. Subsequently, sterile media supplemented with 1 mM glycan, was perfused for 4 hours to initiate phage washout from the mucus layer. Washouts were collected at set time intervals and phages were quantified via subtractive plating on *E. coli* SupE (permissive for both WT Hoc [*am43^-^/44^-^*] and D246N Hoc) and *E. coli* B (only permissive for D246N Hoc). D) Washout of wildtype Hoc and D246N Hoc phages from the gut-on-a-chip under flow with 1 mM of fucosylated glycan 7A (Lacto-N-fucopentaose I) or E) control glycan 5F (α-1,6-mannobiose) over 4 hours. Lines in panels D and E were plotted as mean values with shaded regions representing the standard error of three biological (i.e. chip) replicates per timepoint (N = 3).

With this knowledge, we proceeded to validate the glycan-binding phenotype of D246N Hoc within the mucosal environment. We tested this by competing the D246N Hoc phage against the wildtype Hoc phage in a phage retention and washout assay within the gut-on-a-chip, under the presence of either a fucosylated (Lacto-N-fucopentaose I [7A]) or non-fucosylated (α-1,6-mannobiose [5F]) glycan. We initiated the experiment by infusing three replicate devices, each with a 1:1 ratio of wildtype Hoc and D246N Hoc phages suspended in fucosylated or non-fucosylated glycan solutions, followed by washout of the phages in the same glycan solutions (Fig.4C). We posit that the wildtype Hoc phage, possessing higher affinity to dissolved fucosylated glycans, will be sequestered away from the mucus layer during the initial infusion, while D246N Hoc phage, with its lower affinity to fucosylated glycans, will be selectively retained in the mucus. Consequently, during the washout we expect higher recovery of D246N Hoc phage for extended periods over the wildtype Hoc phage. To allow for subtractive plating, we utilised a wildtype Hoc phage possessing amber mutations on genes *43* and *44* (herein known as T4 am*43^-^/44^-^*) that was permissive only to *E. coli* strain SupE, while the D246N Hoc phage was permissive on both *E. coli* strains B and SupE. Our results show that in the presence of the fucosylated glycan Lacto-N-fucopentaose I [7A], D246N Hoc phage was recovered at higher levels in the first 2 hours of washout and remained detectable up to 4 hours, whereas wildtype Hoc phage was eliminated by ~3 hours (Fig.4D, Supplementary Fig. 5). Conversely, we observed no difference between D246N Hoc and wildtype Hoc phage washout in the presence of a control non-fucosylated glycan, α-1,6-mannobiose [5F], where both phages persisted up to the 4-hour final timepoint (Fig.4E, Supplementary Fig. 5). Overall, we were able to elucidate and verify the phenotypic effect of the D246N Hoc with fucosylated glycans (as predicted via SPR) within the gut-on-a-chip mucosal environment.

## Discussion

Phages are largely considered inert with respect to the mammalian “host” and chiefly respond to antagonistic selection from their immediate replicative bacterial hosts. However, the mammalian milieu – in this case, the gut mucosa – is also a complex environment that can impose additional selection pressures such as mucus turnover dynamics and glycosylation that act on both bacterial and viral entities^28,30,31^. Here, we demonstrated that phages evolve in response to a dynamic mammalian gut mucosal environment, revealing a trans-domain evolution along the phage-mammalian axis. Unlike phage-bacteria antagonistic co-evolution where selection is largely directed and predictable (Fig.2C, test-tube), selection imparted by the mammalian mucosal environment is subtler as evidenced by variations in mutational profiles across evolving phage populations observed in individual gut-on-a-chip replicates (Fig.2C, gut-on-a-chip). The disparity observed in gut-on-a-chip population dynamics is reflective of interpersonal variations seen in gut viral community dynamics^2,3^. We speculate that stochastic ecological effects arising from demographic noise and gut spatial complexity^32^, could be key factors in determining mucosal selection within independent gut environments. Despite the mutational disparity and ecological variation between mucus-evolving phage populations, we acquired a genetically-recombined phage mutant exhibiting altered affinity towards fucosylated glycans (Fig.4C) via a mutation in the phage capsid’s mucus-adhering Hoc domain (Fig.4A). This mutation conferred a fitness advantage within the mucus layer by altering phage Hoc affinity to mucin glycans, specifically by decreasing Hoc affinity to fucosylated glycan structures (Fig.3E & 4B). While diminished phage glycan-binding may appear counterintuitive as a fitness advantage for persistence in the mucosal environment, we note that: i) the exact glycosylation profile and glycan abundance of the gut-on-a-chip mucus layer were unknown and that, ii) our SPR screen was limited to a small subset of fucosylated glycans. Nonetheless, the mutation lent phages a detectable phenotypic response within the mucosal environment (Fig.4D & E); thus, validating the evolutionary interaction between the phage and the mammalian mucus layer.

Mucin fucosylation is widespread along the gastrointestinal tract of functional *FUT2* human genotypes (known as “secretors”), especially within the proximal and distal colon^31^. This suggests that the human host genotype and glycosylation demography directly influences gut phage biogeography at the inter- and intra-individual level, respectively. Moreover, the majority of the gut phageome possesses open reading frames for variable glycan-binding superfamily domains^5,33^ suggesting that gut phages have immense adaptive freedom to respond and co-evolve with an individual’s unique mucosal glycosylation patterns to foster persistence^10^. Successful phage variants that emerge and persist in the gut to achieve high abundances will therefore, have greater capacities for genetic recombination to promote the fixation of beneficial mutations within the population. This subsequently dictates the phage populations that will reside and further engage in co-evolution with both the individual’s gut microbiome and gut environment. Alongside antagonistic co-evolution with gut bacteria, this novel symbiosis between the mammalian gut and phages might lend toward stable, long-term and highly personalised viromes and microbiomes, which are often recapitulated in human metagenomic cohort studies^2,34^. Overall, our findings may have far-reaching implications on re-evaluating phage evolution beyond antagonistic co-evolution with bacteria. In particular, we envisage future directions towards human host-centric intelligent phage design in synergy with host-directed phage evolution for highly personalised medicine and refined *in vivo* phage applications.

## Supporting information

Breseq alignment and mutational comparisons

## Acknowledgements

This work, including the efforts of J.J.B., was funded by the Australian Research Council (ARC) Discovery Early Career Researcher Award (DECRA) (DE170100525). This work was performed in part at the Melbourne Centre for Nanofabrication (MCN) in the Victorian Node of the Australian National Fabrication Facility (ANFF).

## Author contributions

Conceptualisation, resources and funding acquisition were carried out by J.J.B. The work was supervised by M.J.M, A.N. and J.J.B. Experimental design was carried out by W.H.C, M.J.M, A.N. and J.J.B. Evolution experiments (including phage purification, DNA isolation, extraction and bioinformatics), recombination assay, sequencing-based phage competition and competitive phage-glycan washout assay were conducted by W.H.C. Gut-on-a-chip fabrication, culture and set-up were performed by W.H.C, C.K. and C.D. with automated dispensing platform design and realisation by D.M. and Y.Z. High resolution phage sampling experiment was performed by W.H.C with qPCR quantification performed by C.K. Molecular cloning of recombinant Hoc protein expression strains was performed by R.P. and W.H.C. Recombinant Hoc protein expression, purification and modelling were performed by R.B. and T.L. Glycan array and SPR experiments with full data processing and analysis were performed by O.C. and J.T. Formal analysis of results were done by W.H.C, L.W., M.J.M. and J.J.B. The original draft was written by W.H.C. with subsequent reviews by M.J.M. and J.J.B and edits by W.H.C and J.J.B. All authors read and commented on the final draft of the manuscript.

## Methods

### Culture protocol for bacteria, phage and tissue culture cell lines

*Escherichia coli* strain B was used for all experiments and was grown in LB medium (10 g Tryptone, 10 g NaCl, 5 g yeast extract in 1 L of sterile dH2O) at 37°C with agitation. T4 phage, which uses *E. coli* strain B as a replicative host, was used for all experiments except T4 replication-negative *43^-^* (DNA polymerase) and *44^-^* (polymerase clamp holder subunit) i.e. T4 *am43^-^/744^-^* phage, that only uses amber-permissive host *E. coli* SupE to replicate. The cell line used was a human colon-derived tumorigenic goblet cell, HT29-MTX-E12, obtained from the European Collection of Authenticated Cell Cultures and cultured at 37°C with 5% CO2 in complete media: DMEM with 10% FBS, 1× MEM non-essential amino acids and 1× penicillin-streptomycin antibiotics (ThermoFisher Scientific). Terminal cellular differentiation was induced with 10 μM N-[N-(3,5-Difluorophenacetyl)-L-alanyl]-S-phenylglycine t-butyl ester (DAPT; Sigma-Aldrich) while mucus-secretion was enhanced with 10 nM phorbol 12-myristate 13-acetate (PMA; Sigma-Aldrich).

### Fabricating the gut-on-a-chip mould and device

A chip mould with 500 μm wide and 350 μm high channel was designed using SolidWorks^®^ 2017 (Dassault Systèmes). The moulds were then 3D-printed and surface-salinized at Melbourne Centre for Nanofabrication (MCN), Victoria. The chips were manufactured by casting a 10:1 mixture of Sylgard^™^ PDMS and its curing agent respectively (Dowsil, USA), onto the moulds and were cured at 90°C until completely solidified. The chips were then removed, trimmed and their inlet and outlet ports were punched. Subsequently, the chips were washed in pentane and acetone to remove residual uncured PDMS followed by plasma bonding the chip onto a glass slide to enclose the chip channel.

The chip channel was ethanol (80%v/v)-sterilised, UV-sterilised and pre-treated with 1:50 MaxGel^™^ ECM (Sigma-Aldrich). The channel was then seeded with 10 μl of HT29-MTX-E12 cells at 3.0 × 10^5^ cells. The seeded chip was incubated statically for 16 hours to allow cell attachment. This was followed by perfusing the attached cells with complete media for 24 hours at 40 μl/hr flow rate to establish a confluent cell layer. The cell layer was then perfused with antibiotic-free media supplemented with cell-inducers DAPT and PMA, for another 24 hours at 120 μl/hr to purge residual antibiotic-containing media from the channel environment and to promote terminal cellular differentiation and mucus secretion by the cell layer. Perfusion was mediated by a 10-channel syringe pump (KD Scientific, USA).

### High temporal resolution gut-on-a-chip phage-bacteria sampling

An in-house automated dispensing platform was constructed to aid sample collection from the gut-on-a-chip over 24 hours at 30-minute intervals. The platform consisted of conveyer belts connected to 5V motors powered by an Arduino circuit board (Arduino, Italy). Two conveyer belt systems were aligned perpendicular to each other allowing motion along the X-Y plane. A custom-made tube holder was connected to the conveyer belt system that holds the gut-on-a-chip tube over the 96-well plate to facilitate sample dispensing into wells. Time-steps for dispensing at 30-minute intervals were coded into Arduino in C++ using Arduino Integrated Development Environment (IDE). For a user-friendly interface, the code was translated onto a virtual switch board executable program using LabVIEW v.2020 (National Instruments, USA). The temporal experiment is initiated by perfusing the gut-on-a-chip with 10^4^ colony forming units (CFU) of *E. coli* B followed by 10^4^ PFU of T4 phages and the device was allowed to run for 24 hours under a 120 μl/hr flow rate whilst connected to the automated dispensing platform to collect egressing fluid samples. In between the gut-on-a-chip and the dispensing platform, the egressing fluid was channelled through an 80°C-heated blank chip to arrest phage and bacterial replication during their egress from the gut-on-a-chip before dispensing. Phages and bacteria from the heat-inactivated samples were quantified using qPCR using SYBR Green I Master with the Lightcycler^®^ 480 (Roche). qPCR primers and cycling protocols for *E. coli* B were as described^35^ using 1 μl of template. T4 protocols was adapted from^36^ using forward primer: 5’-AGGAGTTATATCAACTGTAA - 3’, and reverse primer: 5’-ATCTAGGATTCTGTACTGTT - 3’, with the following cycling protocol: initial denaturation at 95°C for 5 minutes; 40 cycles at 95°C for 30 seconds, 56°C for 30 seconds, 72°C for 30 seconds; using 1 μl of template.

### Phage experimental evolution in gut-on-a-chip

10^4^ PFU of T4 phages were perfused through the gut-on-a-chip followed by 10^4^ CFU of *E. coli* B to supply the phages with hosts to replicate within the chip. The co-culture in each chip was maintained under a 120 μl/hr flow rate with antibiotic-free media for 24 hours. Subsequently, the mucus and the cell layer were collected via washes with 1 × DPBS and 0.25% Trypsin (ThermoFisher Scientific). The chip sample was centrifuged to obtain the bacterial cell pellet, which was resuspended in 100 μl 1 × DPBS. The supernatant containing the phages was treated with 10% chloroform to obtain a purified phage lysate. Phages and bacteria were enumerated using soft-agar overlay assay and colony spot assay, respectively. For our phage passage protocol, 10^4^ phage PFU were taken from the purified phage (supernatant) lysate to inoculate a new gut-on-a-chip with 10^4^ ancestral *E. coli* B CFU. We adopted this passage protocol for a total of 5 passages. In our control experimental evolution, a shaking test-tube was used in place of the gut-on-a-chip within the flow set-up. The passage protocol in the control experiment was the same as the passages of the gut-on-a-chip phage experimental evolution.

### Phage DNA isolation, purification, sequencing and analyses

To obtain sufficient DNA yield for sequencing, phages from all transfers including the ancestral phage population were amplified to high titres (≥10^9^/ml). The phages were amplified by inoculating 30 μl of phage lysate sample into 3 ml of *E. coli* B bacteria in exponential phase (OD_600_ = 0.3). The inoculum was incubated for a maximum of 4 hours at 37°C with agitation to ensure that all phage genotypes have equal probability in expanding without interference from host-induced bottlenecks at late stage incubations. This was followed by 10% chloroform treatment to purify the amplified phage lysate. Phages were concentrated and ultrapurified following the phage-on-tap protocol^37^. 1 ml of each ultrapurified phage passage lysate was treated with 10 μl Ambion^™^ DNase I (ThermoFisher Scientific) and 20 μl RNase (Sigma-Aldrich) to eliminate bacterial genome contamination. Subsequently, the lysates underwent phage DNA extraction using Phage DNA Isolation Kit (Norgen Biotek^®^, Canada) as per manufacturer protocol with the following modification to maximise DNA yield: 10 μl of 20 mg/ml Proteinase K (Sigma-Aldrich) per 1 ml of amplified phage lysate and incubated at 55°C for 1.5 hours. Phage DNA quality and concentrations were assessed via Nanodrop A_260/280_ (ThermoFisher Scientific) readout and QuBit^®^ Fluorometric Quantification High Sensitivity assay (ThermoFisher Scientific), respectively. Phage DNA samples were sequenced using Illumina HiSeq^®^ 150bp paired-end chemistry (GeneWiz^®^, Hong Kong) and read alignments to T4 reference genome^21^ (NCBI GenBank ID: MT984581.1) were performed via the Breseq Polymorphism Mixed Population pipeline with filter settings turned off to maximise variant calling. *De novo* mutation hits were derived by comparing evolved phage population hits with ancestral background mutations using Breseq’s-gdtools `SUBTRACT` and `COMPARE` commands.

### Lytic phage recombination assay

T4 Δ21bp *goF* mutant was isolated from transfer 4 chip-evolved replicate 1 population by isolating phage plaques from soft-agar overlay. The phage isolates were PCR-screened and Sanger-sequenced with the flanking *goF* primers i.e. forward: 5’ – GCATTAATCAGCATCAGTAC − 3’ and reverse: 5’ – AAGACGGCACAACTTACTGG – 3’, with the following PCR protocol: initial denaturation at 95°C for 10 minutes; 34 cycles at 95°C for 10 seconds, 57°C for 15 seconds, 72°C for 60 seconds; and final elongation at 72°C for 5 minutes. T4 *hoc* knockout (Δ*hoc*) phage was also PCR-amplified and sequence-confirmed using the flanking *hoc* primers i.e. forward: 5’ – GCTGAAACTCCTGATTGGAAATCTCACCC – 3’ and reverse: 5’ – GCCCATAATACAGCCACTTCTTTTGCC – 3’, with the following PCR protocol: initial denaturation at 95°C for 10 minutes; 34 cycles at 95°C for 30 seconds, 60°C for 60 seconds, 72°C for 90 seconds; and final elongation at 72°C for 10 minutes. The verified phages were amplified and chloroform-purified to high titre (≥10^9^ PFU/ml), respectively. The phages were diluted in SM buffer (5.8 g NaCl, 2.0 g MgSO_4_.7H_2_O, 50 ml 1 M Tris-HCl pH 7.4 in 1 L ddH_2_O) to obtain a 1:1 phage mix containing Δ21bp *goF* and *Δhoc* at 1 × 10^9^ PFU/ml. 1 ml of the mixture was reserved as an initial condition control to test for 1:1 mix accuracy. The remaining mixture was used to prepare four experimental set-ups: two replicates of MOI = 10 and two replicates at MOI = 0.1. In MOI 10, 1 ml of the 1 × 10^9^ PFU/ml mixture was added to 1 ml of 1 × 10^8^ CFU/ml *E. coli* B; while in MOI 0.1, the phage mixture was diluted to 1 × 10^7^ PFU/ml before adding to 1 × 10^8^ CFU/ml *E. coli* B. The co-cultures were then incubated at 37°C with 150 rpm agitation for 30 minutes to allow a one-step T4 phage growth curve. The co-cultures were subsequently quenched with 10% chloroform. The phages in co-culture and the reserved initial condition phage mix were plated via soft-agar overlay. Single plaque cores were obtained from well-separated plaques, resuspended in 100 μl SM buffer, and PCR screened for recombinants (double mutant: Δ21bp *goF* + *Δhoc* or WT recombinant T4 genotypes) using flanking *goF* primers and internal *hoc* primers. Internal *hoc* PCR primers were, forward: 5’ - ACATTATCTACGCTCCAAGC – 3’ and reverse: 5’ - ATCTAGGATTCTGTACTGTT - 3’, with the following protocol: 95°C for 10 minutes; 34 cycles at 95°C for 10 seconds, 56°C for 15 seconds, 72°C for 60 seconds; and final elongation at 72°C for 5 minutes. All PCR products were loaded on 2% agarose gel, stained with SYBR^™^ Gold Nucleic Acid Gel Stain (ThermoFisher Scientific), for 30 minutes at 60V and subsequently, 30 minutes at 50V to allow better separation between the WT and Δ21bp *goF* product. Both *goF* and *hoc* PCR products were matched to their sample of origin in the agarose gel run. The frequency of recombinants was quantified based on the *goF* PCR product size and the presence and absence of *hoc* PCR product.

### Sequencing-based phage competition assay

Wildtype T4 phage and experimentally evolved D246N T4 mutant phage were isolated via plaque coring as previously described. The cores were resuspended in 100 μl of SM buffer and samples were PCR-amplified with flanking *hoc* primers i.e. forward: 5’ – GCCCATAATACAGCCACTTCTTTTGCC – 3’ and reverse: 5’ – GCTGAAACTCCTGATTGGAAATCTCACCC – 3’, with the following protocol: initial denaturation at 95°C for 10 minutes; 30 cycles at 95°C for 30 seconds, 60°C for 60 seconds, 72°C for 90 seconds; and final elongation at 72°C for 10 minutes. The verified phages were amplified and chloroform-purified to high titre (≥10^9^ PFU/ml), respectively. The amplified phages were diluted in antibiotic-free tissue culture media to obtain a 1:1 phage mix containing WT and D246N phages at 2 × 10^6^ PFU (1 × 10^6^ PFU each). 1 × 10^6^ PFU of the phage mix was reserved as an initial condition i.e. T = 0 control. Three gut-on-a-chip replicates were each infused with 10^6^ CFU *E. coli* B bacteria followed by 1 × 10^6^ PFU phage mix at 120 μl/hr flow rate. The inoculated devices were maintained at 120 μl/hr for 24 hours and egressing fluid samples were collected for 1 hour at the 24-hour timepoint. Fluid samples were collected in 1 ml SM buffer to rapidly dilute the collected phages and bacteria to limit further phage adsorption during sample collection. Collected samples were then amplified, DNA-extracted, sequenced and analysed as previously outlined to track the frequency of D246N mutant phage as it competes with WT T4 phage over 24 hours. Selection coefficients were calculated as described in Supplementary Table 2 based on absolute reads, obtained by multiplying read depth and coverage, of the mutation.

### Molecular cloning of recombinant Hoc protein expression strains

Wildtype Hoc T4 phage and D246N Hoc T4 phage genomic DNA were extracted as described above. The respective *hoc* genes were PCR-amplified using primers designed with Ncol/Spel restriction sites i.e. forward: 5’ – CCTCCATGGCGATGACTTTTACAGTTGATATAAC – 3’ and reverse: 5’ – TTGACTAGTTATGGATAGGTATAGATGATAC – 3’, with the following protocol: initial denaturation at 98°C for 5 minutes; 36 cycles at 98°C for 30 seconds, 58°C for 30 seconds, 72°C for 120 seconds; and final elongation at 72°C for 5 minutes. The amplified *hoc* products were gel-extracted following manufacturer’s protocol (GenElute^™^ Gel Extraction Kit, Sigma Aldrich). Wildtype and D246N *hoc* genes were individually cloned in-frame to expression vector pPROEX-HTb, containing an N-terminal hexa-His sequence. Briefly, the amplified *hoc* product and pPROEX-HTb were digested with NcoI and SpeI (New England Biolabs) at 37°C overnight, followed by ligation at room temperature for 2 hours. The ligated expression vector was transformed into NEB 5α Competent *E. coli* as per manufacturer’s protocol (New England Biolabs) and plated on LB medium supplemented with 100 μg/ml ampicillin, where colonies were PCR-screened as above mentioned. PCR-positive colonies were grown and the vector was extracted using GenElute Plasmid Miniprep Kit following manufacturer’s protocol (Sigma-Aldrich). The vector was then transformed into expression strain *E. coli* BL21(DE) Star as follows. *E. coli* BL21(DE) Star was grown in LB medium to OD_600_ 0.4 at 37°C. 5 ml of culture was centrifuged at 4°C and the pellet was washed thrice with 1 ml ice-cold 10% glycerol between centrifugations. The pellet was resuspended in 50 μl of ice-cold 10% glycerol and added with 3 μl of the expression vector. The mixture was transferred into a 0.1 cm electroporation cuvette (BioRad) and pulsed at 1.8 kV. Electroporated cells were recovered in 1 ml pre-warmed LB medium for 1 hour at 37°C and subsequently plated on LB medium supplemented with 100 μg/ml ampicillin to recover Hoc expression strains.

### Recombinant Hoc protein expression, purification and modelling

Hoc expression strains were grown in Terrific Broth (with shaking) to OD_600_ 0.8 at 37°C. Expression was induced with 0.2 mM IPTG, incubation temperature dropped to 18°C and cells collected by centrifugation the following morning. Cell pellets were resuspended in 20 mM Tris pH8, 300 mM NaCl, 20 mM imidazole, 0.5 mM MgCl_2_, 1× complete EDTA-free protease inhibitor (Roche) and lysed through an Avestin Emulsiflex C3 cell press. Following centrifugation at 18000 ×g the soluble fraction was applied to a 5 ml HisTrap HP column (GE Healthcare). The column was washed and protein eluted along a gradient using 20 mM Tris pH8, 400 mM NaCl, 1 M imidazole. The peak fraction (eluting at ~150 mM imidazole) was pooled and further purified over size exclusion chromatography on a HiLoad 16/600 Superdex 200 pg column (GE Healthcare) equilibrated in SEC buffer (20 mM Tris pH 8.0 and 150 mM NaCl). The Hoc proteins each eluted as a single monomeric peak and were run on reducing SDS-PAGE and verified by anti-His western (R&D Systems) (Supplementary Fig.3). Proteins were concentrated to 1 mg/ml, EDTA added to 0.5 mM final and aliquots snap-frozen in liquid nitrogen. The structural model of the T4 Hoc protein was generated using Phyre2 server^38^ and modelled upon the crystal structure of the three N-terminal IgG domains of phage RB49 Hoc protein (PDB ID: 3SHS). The capsid binding domain could not be accurately modelled due to a lack of solved structural homologues.

### Glycan array printing

Glycan arrays consisting of 150 diverse glycans (DextraLabs) in the absence of spacers were taken from existing glycan libraries^39,41^. Glycans were amine functionalized as previously described^42^ and subsequently printed as described^43^. Briefly, glycosylamines were suspended in 1:1 dimethylformamide (DMF): dimethyl sulfoxide (DMSO) at a concentration of 500 μM and printed onto SuperEpoxy3 glass slides (ArrayIt) using a SpotBot Extreme array spotter (ArrayIT) in a six-pin subarray print per glass slide format. All glycans were printed in replicates of four, including four AlexaFlour 555/647 and FITC control spots, per subarray using 946MP4 pins and a contact time of 1 second at 50% relative humidity, with pins being reloaded after every 8 spots. DMF: DMSO was also printed as blanks controls. The printed arrays were subsequently acetylated in 25% (v/v) acetic anhydride in methanol at 4°C for 15 min, and then neutralized in 1:1 ethanolamine: DMF. Finally, glycan arrays were washed with 100% ethanol and dried in an empty 50 mL tube by centrifugation for 5 min at 200 ×g. Glycan arrays were vacuum sealed and stored at 4°C.

### T4 phage labelling and glycan array hybridization

To label T4 phages (wildtype [WT], D246N Hoc or *Δhoc*), stocks were diluted to 10^8^ phages/mL in SM buffer and allowed to incubate with SYBR green dye (1:1000) (Molecular Probes) in the dark at 4°C for 1 hour. Excess dye was removed by three consecutive washes with 1 mL of SM buffer using an Amicon ultrafiltration tube (100 kDa). A buffer-exchange through three consecutive washes with 1 mL of array phosphate buffered saline (PBS) (50 mM PBS, 1.8 mM MgCl_2_ and 1.8 mM CaCl_2_, pH 7.4) was similarly performed using Amicon ultrafiltration tubes (100 kDa) (Merck). SYBR-labelled phages were prepared fresh daily and immediately applied to glycan arrays after buffer-exchange. Before hybridizations, glycan array slides were blocked in 0.5% BSA in array PBS for 5 min at room-temperature (RT). After washing with array PBS, slides were dried through centrifugation and a Gene Frame (1.7 × 2.8 cm, 125 μL, Abgene) was used to isolate the arrays prior to the addition of the labelled phage. 10^8^ of either SYBR labelled WT, D246N Hoc or *Δhoc* T4 phages were applied to individual glycan arrays as a 1 mL bubble and allowed to hybridize at RT for 1 hour in the dark. In the final 5 minutes of incubation, phages were fixed through the addition of formaldehyde into the same bubble (final concentration 4%). Following hybridization, glycans arrays were gently washed three times for 5 min in array PBS and finally dried through centrifugation.

### WT and D246N Hoc protein labelling and glycan array hybridization

Labelling of recombinant WT and D246N Hoc proteins was performed using their respective hexa-His-tags. Here, 1 μg of each protein was incubated at a molar ratio of 1:2:4 with anti-His-tag mouse monoclonal antibody (Cell Signalling Technology), anti-mouse-IgG-Alexa647 conjugated rabbit polyclonal antibody (Life Technologies) and goat conjugated anti-rabbit-IgG-Alexa647 polyclonal antibody (Life Technologies) in 1 mL Array PBS. This complex was allowed to hybridize in the dark at 4°C for 15 min. As described previously, gene frames were used to isolate glycan arrays, and Alexa647 labelled recombinant Hoc proteins were applied as a bubble for 1 hour at RT and allowed to hybridize. Glycan arrays were subsequently washed for three times for 5 min in array PBS, and dried through centrifugation.

### Fluorescent image acquisition and data processing

Fluorescence intensities of the array spots were measured with the Innoscan 1100AL (Innopsys) scanner using either the 488 nm (SYBR) or 635 nm (A647) laser excitation wavelength depending on the sample. The Image analysis was carried out using the inbuilt imaging software, MAPIX (Innopsys). Raw glycan signals were exported into Microsoft Excel 2016. The mean background was calculated from the average of DMF/DMSO blanks on the array plus three standard deviations. This was subtracted from each glycan to generate an adjusted signal. A one tailed t-test was performed with significance set at *p* = 0.05. Binding events confirmed across 3 arrays were compiled as heatmaps representing T-test and fold increases above background.

### Surface plasmon resonance detection

Surface plasmon resonance (SPR) experiments to confirm glycan hits and elucidate differences in binding affinity between the WT and D246N Hoc proteins were performed using a Pioneer FE SPR system (Pioneer). WT and D246N Hoc proteins were loaded onto channels 1 and 2 of a HisCap biosensor (Satorious) and channel 3 was blank immobilized to enable reference subtraction in PBS. A minimum of 5000 relative units (RU) of either Hoc protein was immobilized using the nitrilotriacetic acid (NTA)-Nickel capture system modified from reference^44^. Here, the hexa-His-tag allows capture of the Hoc proteins in the correct orientation and subsequent covalent crosslinking prevents protein from dissociating over the course of the SPR run. In brief, nickel was bound to the HisCap biosensor using NiSO_4_ in running buffer. The carboxymethylated dextran (CMD) surface was then activated using N-hydroxysuccinimide (NHS)/1-ethyl-3(3-dimethylaminopropyl)-carbodiimide hydrochloride (EDC). Each protein was subsequently immobilized at flow rate of 10 μL/min for 10 min. Uncoupled amine reactive sites of the CMD were blocked through an injection of ethanolamine and finally 0.35 M EDTA was injected to remove any poorly associated protein. A maximum concentration of 100 μM of selected glycans was tested using a OneStep analysis programmed using the Pioneer instrument. OneStep was performed with 75% loop volume and a 3% sucrose control. Glycans were flowed at 40 μL/min with a dissociation time of 180s (Supplementary Fig.5). Subsequent regeneration of the surface was performed with TE buffer for 60s at 50 μL/min and 60s dissociation. Blanks were run periodically every 2 cycles. Analysis of SPR sensorgrams to determine glycan dissociation constants (*K_D_*) was performed separately with the Qdat analysis software package (Biologic Software, Campbell, Australia). All analyses were performed on two independently prepared HisCap chips with each protein loaded twice and glycans tested in duplicate per run. SPR responses less than 5 RU were deemed insignificant and attributed to non-specific interaction of the glycan with the positively charged HisCap chip surface.

### Phage retention and washout assay

A 1:1 phage mix consisting of D246N Hoc T4 phage and a WT T4 Hoc phage (*am43^-^/44^-^*) was prepared in antibiotic-free tissue culture media at 1 mM final glycan concentration of glycans α-1,6-mannobiose (DextraLabs) or Lacto-N-fucopentaose I (DextraLabs). The 1:1 phage ratio was verified by plating on *E. coli* SupE and *E. coli* B lawns in triplicates where, the amber phage only plaques on an amber mutant permissive host, *E. coli* SupE while D246N Hoc phage plaques on both *E. coli* SupE and *E. coli* B. Hence, we were able to quantify the D246N Hoc phage (on *E. coli* B) and the amber mutant phage via subtraction (total plaques from *E. coli* SupE – total plaques from *E. coli* B). Three replicate gut-on-a-chips were infused with 1 *×* 10^7^ PFU/ml of 1:1 phage-glycan mix for 1 hour at 120 μl/hr. After which, the devices were perfused with sterile antibiotic-free tissue culture media supplemented with 1 mM of the appropriate glycan for 4 hours. Device effluents were collected in equal volumes of SM buffer every 15 minutes for the first hour and every 30 minutes for the subsequent hours. The phage timepoints were quantified by spot-plating the device effluents on both *E. coli* B and *E. coli* SupE lawns to assess for phage washout.

