## Supplementary information for "Bacteriophage adaptation to a mammalian mucosa reveals a trans-domain evolutionary axis"

**Supplementary Figure 1**

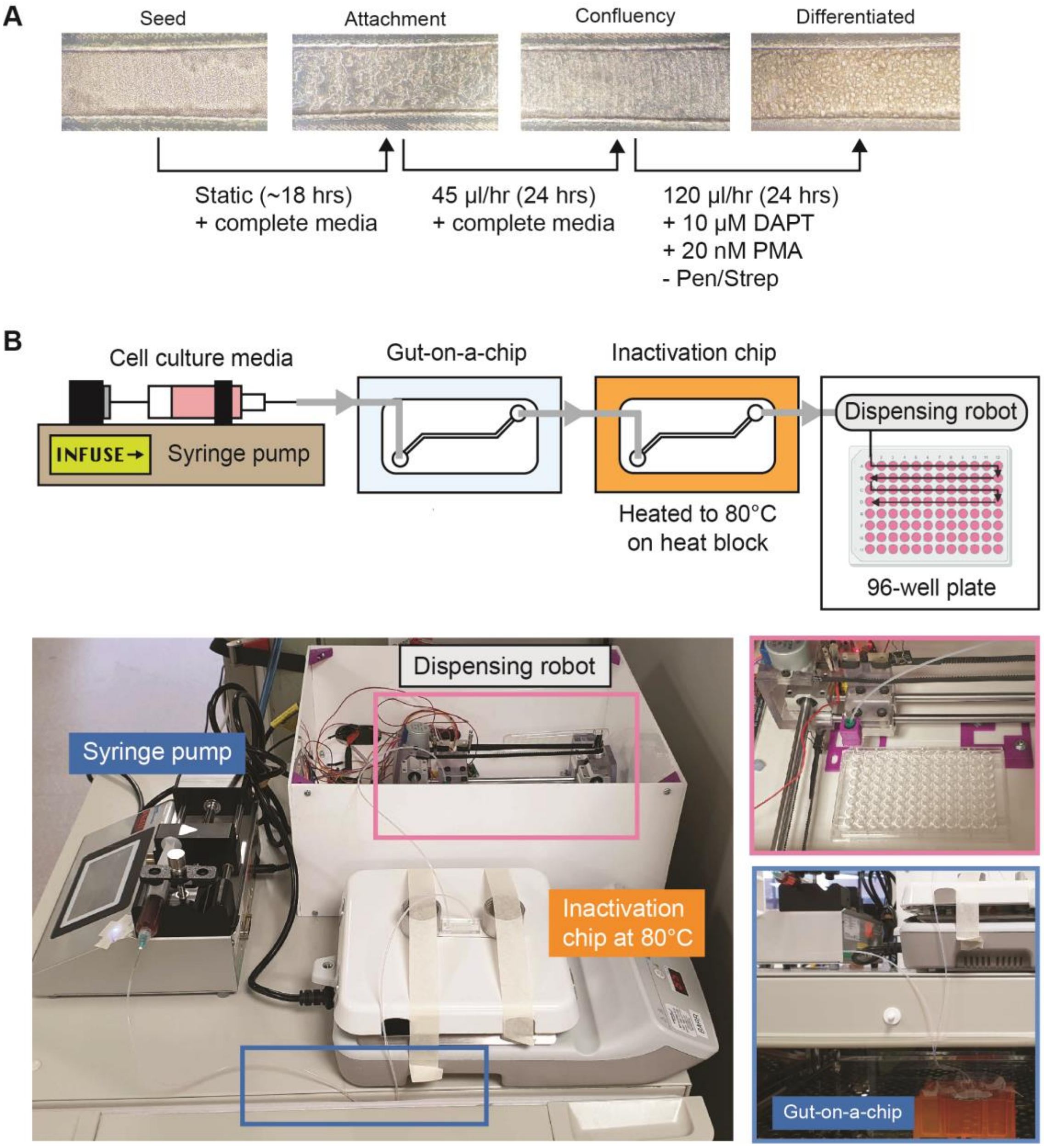


**Supplementary Fig.1:** A) HT29-MTX-E12 cell culture in the gut-on-a-chip. Cells were seeded to a density of $3.0 \times 10^7$ cells/ml in the device channel and allowed to incubate under static conditions for ~18 hours for attachment. Attached cells were subsequently fed with tissue culture media at low perfusion rate of 45 µl/hr until the cell layer has established confluency (~24 hours). Following that, antibiotic-free (-Pen/Strep) media, supplemented with inducers - 10 µM N-[N-(3,5-Difluorophenacetyl)-L-alanyl]-S-phenylglycine t-butyl ester (DAPT) and 20 nM 12-myristate 13-acetate (PMA) – to promote terminal cellular differentiation and mucus production, respectively were perfused at a 120 µl/hr to mimic physiological shear stresses of the *in vivo* gut. The resulting acclimatised cell layer possessed extensive goblet cell morphologies and exhibited mucus production visible from alterations in cell layer shade and opacity under light microscopy. B) Automated sampling platform for high temporal resolution sampling from the gut-on-a-chip. The gut-on-a-chip was perfused by a syringe pump where egressing fluid from the device was channelled to an “inactivation chip” – an empty device placed on a heat-block at 80°C – to inactivate phages and bacteria within the egressing fluid sample. The inactivated sample was subsequently channelled to a dispensing robot which dispenses the inactivated sample across a sterile 96-well plate over 24 hours, which were then quantified for phage and bacteria abundances via qPCR. The robot was manufactured in-house, consisting of Arduino-driven stepper motors and conveyer belts that moves along the horizontal X-Y plane following written code with relevant time-steps.

### Supplementary Figure 2

**A.** *goF* nucleotide sequence:

```
Mutant  1    ATGGCTATTAAATTTGAAGTTAATAAATGGTATCAATTTAAAAATAAACAAGCTCAAGAA  60
             ||||||||||||||||||||||||||||||||||||||||||||||||||||||||||||
Ancest  1    ATGGCTATTAAATTTGAAGTTAATAAATGGTATCAATTTAAAAATAAACAAGCTCAAGAA  60

                   1)Sub:T
Mutant 61    AATTTTATTAAAGACCATACTGATAACGGAATCTATGCACGACGTTTAGGTATGGAGCCT  120
             ||||||||||||||||||||||||||||||||||||||||||||||||||||||||||||
Ancest 61    AATTTTATTAAAGACCATACTGATAACGGAATCTATGCACGACGTTTAGGTATGGAGCCT  120

                         2)5aa-del
Mutant 121   TTTAAAATTTTAGATGCTGATTATCTTGGGCGTCCTACTAAAATTATGACATCCATAGGT  180
             ||||||||||||||||||||||||||||||||||||||||||||||||||||||||||||
Ancest 121   TTTAAAATTTTAGATGCTGATTATCTTGGGCGTCCTACTAAAATTATGACATCCATAGGT  180

Mutant 181   GTACTCAAACGTTGTGCCGGCGGTGATATCCTTGACGAAAACTTTATCTGGCTCTCTACT  240
             ||||||||||||||||||||||||||||||||||||||||||||||||||||||||||||
Ancest 181   GTACTCAAACGTTGTGCCGGCGGTGATATCCTTGACGAAAACTTTATCTGGCTCTCTACT  240


                3)Sub:A                                          7aa-del
Mutant 241   AACGAAGCTGGGTTCTTTGATGAAGTGGAAAATCCATATCAGGCGGTTG-----------  289
             |||||||||||||||||||||||||||||||||||||||||||||||||
Ancest 241   AACGAAGCTGGGTTCTTTGATGAAGTGGAAAATCCATATCAGGCGGTTGAAGAGCAAGAG  300

Mutant 290   ----------AAGAGCAAGAACAAATAGAAGATTTCACAGAATTCCCAGTCATGAAAGTT  339
                       ||||||||||||||||||||||||||||||||||||||||||||||||||
Ancest 301   CAGGAAGAGAAAGAGCAAGAACAAATAGAAGATTTCACAGAATTCCCAGTCATGAAAGTT  360
      4)Sub:T (early STOP codon: UAG)

Mutant 340   ACTATTGAAAATAATGATCAGGCGTGGTCTTTATATCAGATGTTGAAAGCTTACTTTAAG  399
             ||||||||||||||||||||||||||||||||||||||||||||||||||||||||||||
Ancest 361   ACTATTGAAAATAATGATCAGGCGTGGTCTTTATATCAGATGTTGAAAGCTTACTTTAAG  420

Mutant 400   GAATAA  405
             ||||||
Ancest 421   GAATAA  426
```

**B.** GoF protein sequence:

```
Mutant 1     MAIKFEVNKWYQFKNKQAQENFIKDHTDNGIYARRLGMEPFKILDADYLGRPTKIMTSIG  60
             ||||||||||||||||||||||||||||||||||||||||||||||||||||||||||||
Ancest 1     MAIKFEVNKWYQFKNKQAQENFIKDHTDNGIYARRLGMEPFKILDADYLGRPTKIMTSIG  60

Mutant 61    VLKRCAGGDILDENFIWLSTNEAGFFDEVENPYQAVE-------EQEQIEDFTEFPVMKV  113
             |||||||||||||||||||||||||||||||||||||       ||||||||||||||||
Ancest 61    VLKRCAGGDILDENFIWLSTNEAGFFDEVENPYQAVEEQEQEEKEQEQIEDFTEFPVMKV  120

Mutant 114   TIENNDQAWSLYQMLKAYFKE  134
             |||||||||||||||||||||
Ancest 121   TIENNDQAWSLYQMLKAYFKE  141
```

**Supplementary Fig.2:** A) Nucleotide sequence of complete (ancestral; `Ancest`) *goF* gene from the National Centre for Biotechnology Information (NCBI) aligned with Sangersequenced *goF* mutant (Mutant). The alignment shows the position of the in-frame 21bp-deletion (Δ21bp) in yellow. Other previously characterised mutations are shown in blue, numbered and labelled in bold[22]. B) The corresponding translated protein sequence of the ancestral *goF* gene aligned with the mutant *goF* protein sequence, demonstrating the seven amino acids eliminated by the in-frame Δ21bp in yellow. The green zones represent the acidic region with predicted homology to RNA-binding proteins and RNA helicases[45].

**Supplementary Figure 3**

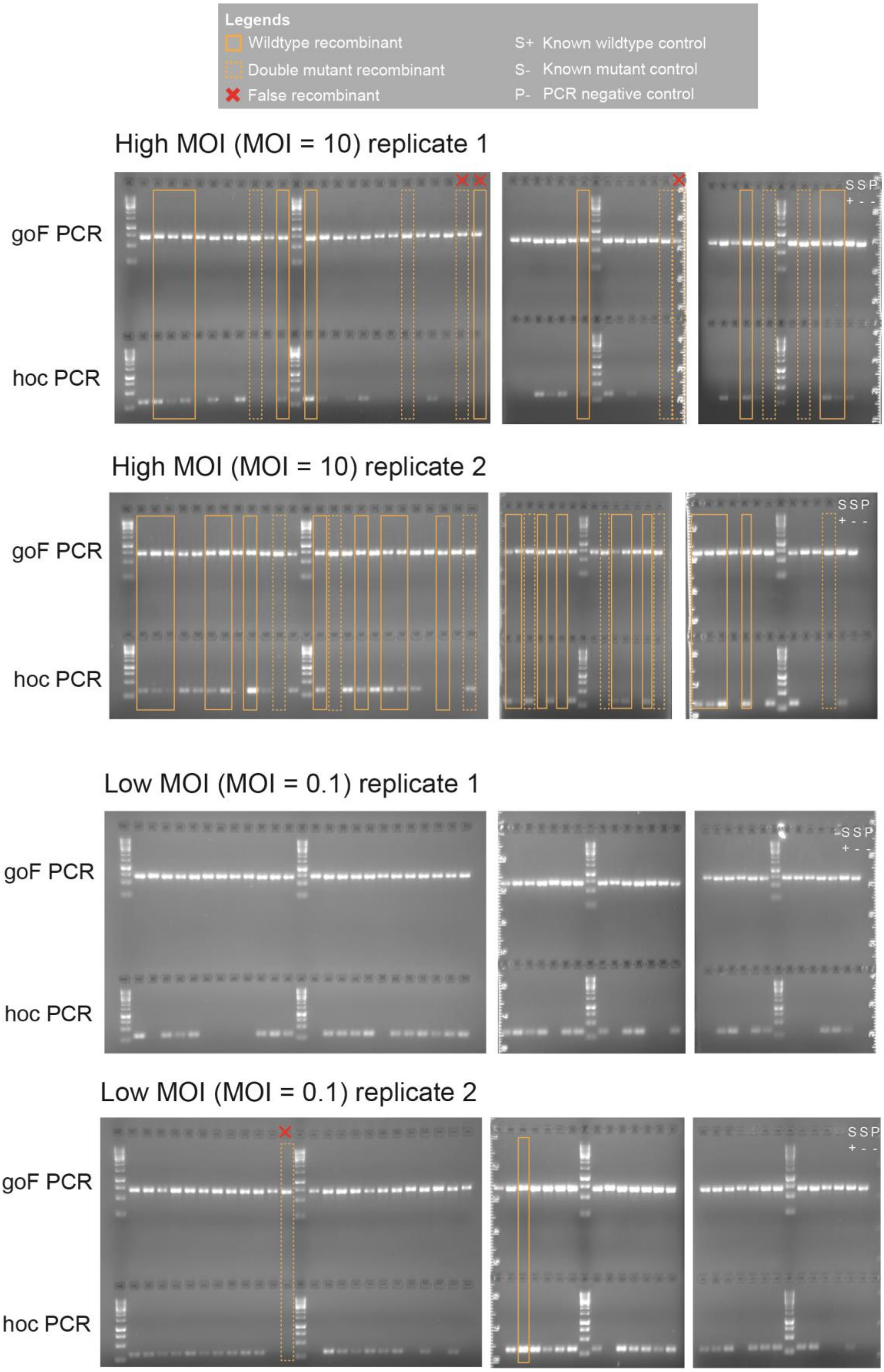

**Supplementary Fig.3:** PCR gels of phage recombinant screenings from recombination experiment at high and low multiplicities-of-infections (MOIs 10 and 0.1), with two independent replicates per MOI condition. A total of 49 phage isolates were screened per condition per replicate alongside three controls: i) known wildtype sample control (S+), known mutant control (S-), and sample negative PCR control (P-) to assess for contamination. PCR for *goF* on the top row of the gels discriminates between wildtype (ancestral) and the mutant phage genotypes by observing for band shifts corresponding to the $\Delta$21bp *goF* mutation. Meanwhile, PCR for *hoc* on the bottom row of the gels discriminates between wildtype and $\Delta$*hoc* phage genotypes for the presence or absence of PCR product, respectively. Solid rectangles indicate wildtype recombinants, possessing both wildtype alleles of the *goF* and *hoc* gene, while dashed rectangles indicate double mutant recombinants, possessing both $\Delta$21bp *goF* and $\Delta$*hoc* mutations. All recombinants were verified via a separate PCR and false recombinants were labelled accordingly (i.e. red crosses) and were discounted from the total recombinant count. We note that the false recombinants were largely arising from ambiguous amplified products in the initial PCR screen.

**Supplementary Figure 4**

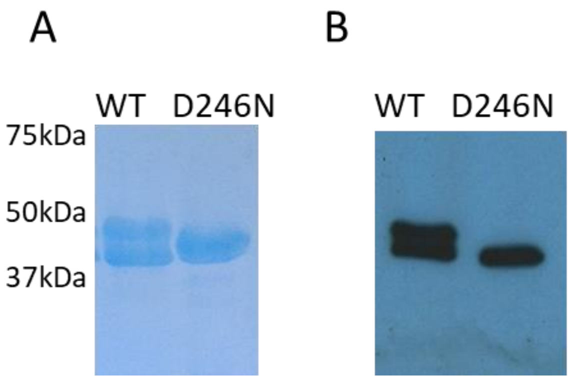


**Supplementary Fig.4**: A) Coomassie stained SDS-PAGE of purified His-WT Hoc and His-D246N Hoc. WT Hoc purified as a doublet. B) Subsequent probing of Western transfer with anti-His antibodies (R&D Systems) shows that both the full-length and partially degraded WT Hoc are His-tagged, implying C-terminal degradation of the protein. The D246N Hoc mutant has also undergone truncation from the C-terminus to produce a stable N-terminally His-tagged Hoc protein of the same size as the WT degradation product. These events did not impact upon the three N-terminal Ig-like domains, with the loss of approximately 5kDa from the C-terminus of the fourth domain (the capsid binding domain).

**Supplementary Figure 5**

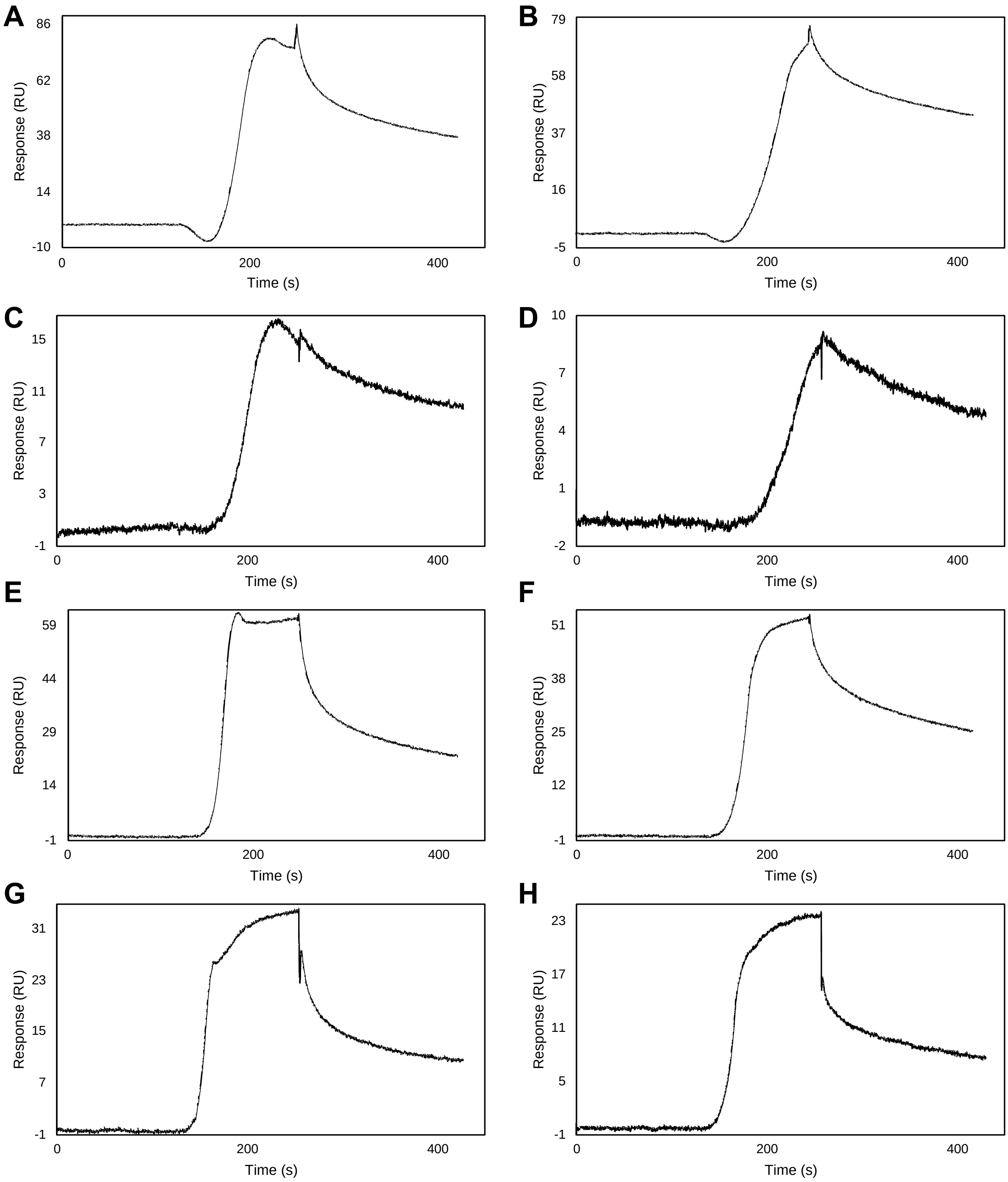


**Supplementary Fig.5:** SPR sensorgrams of the WT Hoc (left column) and D246N Hoc mutant (right column) with various glycans using OneStep analysis. (A-B) 2'-Fucosyllactose; (C-D) Lacto-N-difucohexaose II; (E-F) Lewis$^{y}$, and (G-H) Lacto-N-fucopentaose II.

**Supplementary Figure 5 (*continued*)**

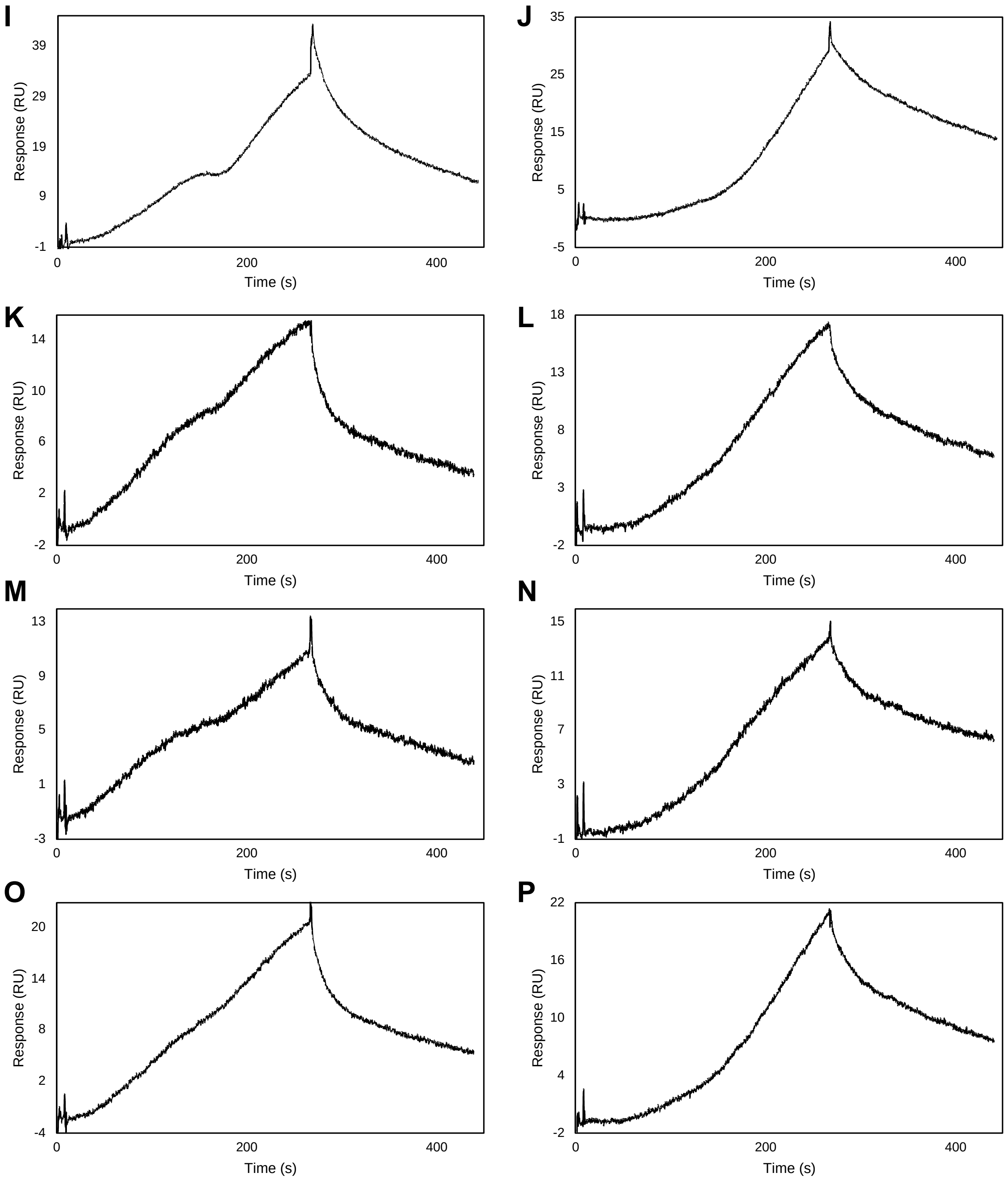


**Supplementary Fig.5 (*continued*):** SPR sensorgrams of the WT Hoc (left column) and D246N Hoc mutant (right column) with various glycans using OneStep analysis. (I-J) Lacto-N-fucopentaose I; (K-L) Lewis$^{a}$; (M-N) Blood Group A Trisaccharide, and (O-P) Lewis$^{x}$.

**Supplementary Figure 6**

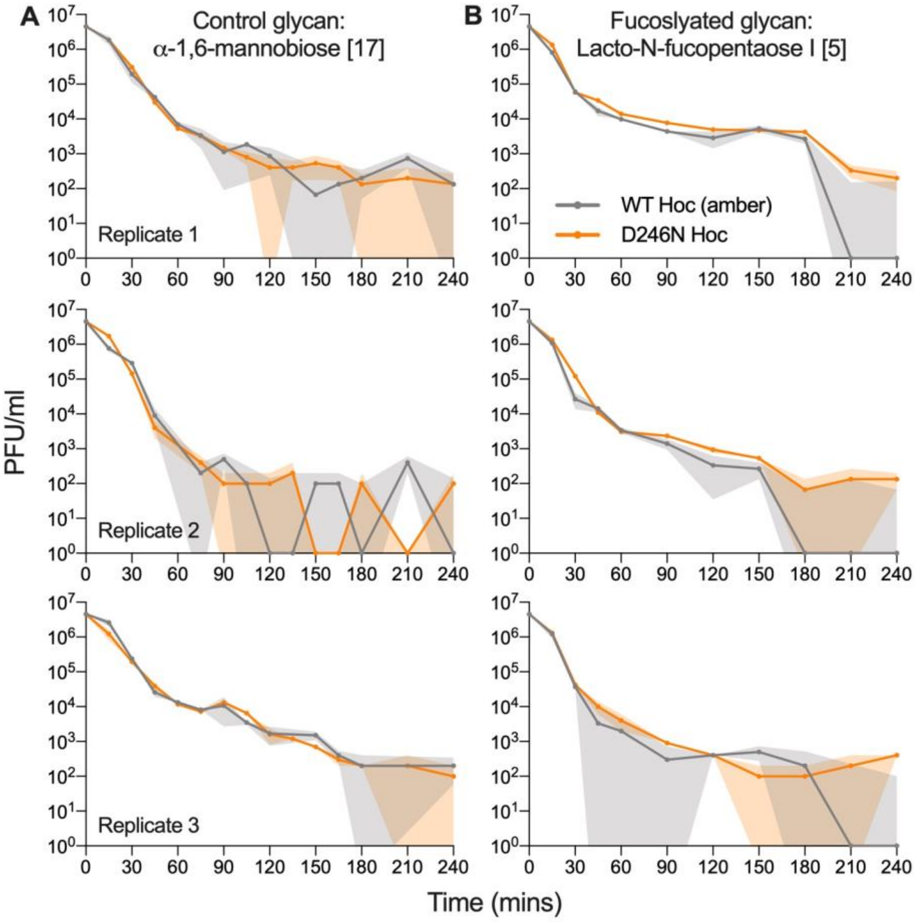


**Supplementary Fig.6:** Individual replicate data for competitive phage-glycan washout from the gut-on-a-chip between WT Hoc (amber *43⁻/44⁻* mutant) and experimentally evolved D246N Hoc mutant under 1 mM A) control glycan: α-1,6-mannobiose and B) fucosylated glycan: Lacto-N-fucopentaose I treatments. Lines are plotted as mean values while shaded regions are standard error of three technical replicates per timepoint (n = 3).

### Supplementary tables

**Supplementary table 1A: List of fixed ancestral and standing background mutations**

Refer to: SI_Table 1A_background_muts.xlsx

**Supplementary table 1B: List of *de novo* mutations.**

Refer to: SI_Table 1B_denovo_muts.xlsx

**Supplementary table 2: Selection coefficient calculations**

Refer to: SI_Table 2_Selection_coefficient_analysis.xlsx

**Supplementary table 3: Whole phage and Hoc protein glycan array heatmaps.**

Refer to: SI_Table_3_glycan array.xlsx

**Scripts (Breseq and mutational comparisons)**

Refer to: SI_alignment_comparison_script.txt
