## Supplementary material for "Bacteriophage adaptation to a mammalian mucosa reveals a trans-domain evolutionary axis": Breseq alignment and mutational comparisons

#MUTATIONAL ALIGNMENT (BRESEQ)

#load Breseq external dependencies (bowtie2 and R)
module load bowtie2/2.3.5 R/3.5.1 breseq/0.33.2

#run script
#replicate = j #transfer = i

#aligning ancestral (T0) population
#running for T0 only
        breseq -j 8 -p --polymorphism-reject-indel-homopolymer-length 0 --polymorphism-reject-surrounding-homopolymer-length 0 --polymorphism-$
        -o T0_nofilter -r T4_sequence.gb T0_L3_1.fq.gz T0_L3_2.fq.gz

#chip replicates
for j in 1 2 3
do
for i in 1 2 3 4 5
        do echo ANALYSING T"$i"GC"$j"_L3_1.fq.gz T"$i"GC"$j"_L3_2.fq.gz
        breseq -j 8 -p --polymorphism-reject-indel-homopolymer-length 0 --polymorphism-reject-surrounding-homopolymer-length 0 --polymorphism-$
        -o T"$i"GC"$j"_nofilter -r T4_sequence.gb T"$i"GC"$j"_L3_1.fq.gz T"$i"GC"$j"_L3_2.fq.gz
done
done

#test-tube control replicates
for j in 1 2 3
do
for i in 1 2 3 4 5
        do echo ANALYSING T"$i"CONT"$j"_L2_1.fq.gz T"$i"CONT"$j"_L2_2.fq.gz
        breseq -j 8 -p --polymorphism-reject-indel-homopolymer-length 0 --polymorphism-reject-surrounding-homopolymer-length 0 --polymorphism-$
        -o T"$i"CONT"$j"_nofilter -r T4_sequence.gb T"$i"CONT"$j"_L2_1.fq.gz T"$i"CONT"$j"_L2_2.fq.gz
done
done


#MUTATIONAL COMPARISON (GDTOOLS)

#Moving .gd files from Breseq output folders to new directory
for j in 1 2 3
do
for i in 1 2 3 4 5
do 
#renaming gd files to correspond with replicate and transfer
mv 'file directory'/output/output.gd 'file directory'/output/T"$i"'exp_conditions'"$j".gd 
#copying renamed files to new directory
cp 'file directory'/output/T"$i"'exp_conditions'"$j".gd 'new file directory'/gdfiles_normalised
done
done

#Subtracting background GC data (removing T0 mutations)
for j in 1 2 3
do
for i in 1 2 3 4 5
do 
gdtools SUBTRACT -o T"$i"'exp_conditions'"$j"_subtracted.gd T"$i"'exp_conditions'"$j".gd T0.gd
done
done

#Comparing subtracted GC table
for i in 1 2 3
do
gdtools COMPARE -o compare_'exp_conditions'"$i".html -r T4_sequence.gb T*'exp_conditions'"$i"_subtracted.gd
done
